# Widely used CaMKII regulatory segment mutations cause tight actinin binding and dendritic spine enlargement in unstimulated neurons

**DOI:** 10.1101/2025.03.18.643923

**Authors:** Ashton J. Curtis, Jian Zhu, Dorota Studniarczyk, Timothy W. Church, Mark Farrant, Matthew G. Gold

**Author notes:** These authors contributed equally (Ashton J. Curtis, Jian Zhu). **Author Contributions**: J.Z., A.J.C, M.F. and M.G. designed research; J.Z., A.J.C., D.S., T.C. and M.G. performed research; J.Z., A.J.C., D.S., M.F. and M.G. analyzed data; M.G. wrote the paper with input from all of the authors. **Competing Interest Statement**: The authors declare no competing interest.

## Abstract

Ca^2+^/calmodulin-dependent protein kinase II (CaMKII) is essential for long-term potentiation (LTP) of excitatory synapses that underlies learning. CaMKII responds to Ca^2+^ influx into postsynaptic spines by phosphorylating proteins and forming new protein interactions. The relative importance of these enzymatic and structural functions is debated. LTP induction triggers CaMKII docking to NMDA receptors, and recent evidence suggests that LTP can proceed without kinase activity after this event. Furthermore, CaMKII interaction with α-actinin-2 is required for dendritic spine enlargement following LTP induction. CaMKII can auto-phosphorylate at T286, which enables autonomous activity after Ca^2+^/CaM dissociation. CaMKII also bears threonine at positions 305 and 306 in its regulatory segment. Experiments with CaMKII variants including a T305A/T306A (‘AA’) double substitution have led to a model whereby T305/T306 phosphorylation by autonomously active CaMKII prevents further Ca^2+^/CaM activation. However, this mechanism is not compatible with some existing data including CaMKII phospho-proteomics and measurements with reporters of CaMKII conformation. Furthermore, autonomous CaMKII activity is now thought to only endure for seconds after LTP induction. In this study, we show that the AA substitution has an unintended gain-of-function property – it enables tight binding to α-actinin-2 in unstimulated neurons. In situ labelling shows that the AA substitution elevates CaMKII-actinin interactions in neurons to a level only normally observed after induction of LTP. The AA CaMKII variant also elevates the proportion of enlarged spines in unstimulated neurons without altering synaptic currents. Calorimetric measurements with purified protein confirm that α-actinin-2 binds tightly to the AA variant of CaMKIIα with no requirement for kinase activation. Using x-ray crystallography, we show that the AA substitution enables α-actinin-2 to adopt a different tighter binding mode. Our findings reinforce the notion that CaMKII primarily fulfils a structural role in LTP.

## Introduction

Changes in the strength of synaptic connections are a fundamental mechanism for encoding new memories and behavioural adaptations (1). In excitatory glutamatergic synapses, large influxes of Ca^2+^ through postsynaptic NMDA receptors (NMDARs) trigger long-lasting changes in the size and responsiveness of dendritic spines known as long-term potentiation (LTP) (2). Ca^2+^/calmodulin (CaM)-dependent protein kinase II (CaMKII) is essential for sensing Ca^2+^ influx (3). CaMKII fulfils both enzymatic and structural roles in LTP: in response to Ca^2+^/CaM activation it phosphorylates receptors and structural proteins, and forms new protein-protein interactions that result in the full expression of LTP including reorganisation of the actin cytoskeleton into enlarged mushroom-shaped spines (3). Historically, the assumption has been that CaMKII phosphorylation is fundamental to LTP but recent studies suggest that its ability to form protein-protein interactions may be more critical (4).

Many studies have focused on understanding how CaMKII activity is regulated by phosphorylation at three sites: T286, T305 and T306 (**Fig. 1***A*). T286 has attracted particular attention since inter-subunit phosphorylation at this site generates a form of the enzyme that retains some activity after Ca^2+^/CaM has dissociated (5–7). Generation of autonomously active CaMKII via T286 phosphorylation was once considered as a means to maintain long-lasting molecular memories of synaptic activation (8). However, more recent work indicates that such phosphorylation only endures for seconds after the Ca^2+^ impulse has receded with a function limited to detecting specific patterns of activation in the initial induction of LTP (9, 10). CaMKII also bears threonine residues at positions 305 and 306 within its regulatory segment (**Fig. 1***A*), and it is widely thought that autonomously active CaMKII can phosphorylate these sites to prevent subsequent activation by Ca^2+^/CaM (11). Many studies have employed CaMKIIα mutants containing the double alanine substitution T305A/T306A (referred to as ‘AA’ hereafter), with the assumption that this mutant reveals the effects of preventing inhibitory phosphorylation. The AA substitution greatly slows the rate of CaMKIIα dissociation from synapses (12). The AA substitution is often combined with a T286D substitution – expression of this CaMKII triple mutant generates highly potentiated synapses that are not amenable to LTD (13–15). However, quantitative proteomics indicates that CaMKII cannot efficiently phosphorylate T305 and T306 (16), and imaging with reporters of CaMKII conformation indicates that inclusion of the T286D alone does not markedly affect the ability of Ca^2+^/CaM to activate the enzyme (9). A possible explanation is that the CaMKII regulatory segment is also important for mediating interactions with α-actinin-2 that underlie spine remodelling (17, 18). Accordingly, in this study we determined the effect of the AA substitution on this aspect of CaMKII function.

**Figure 1.**
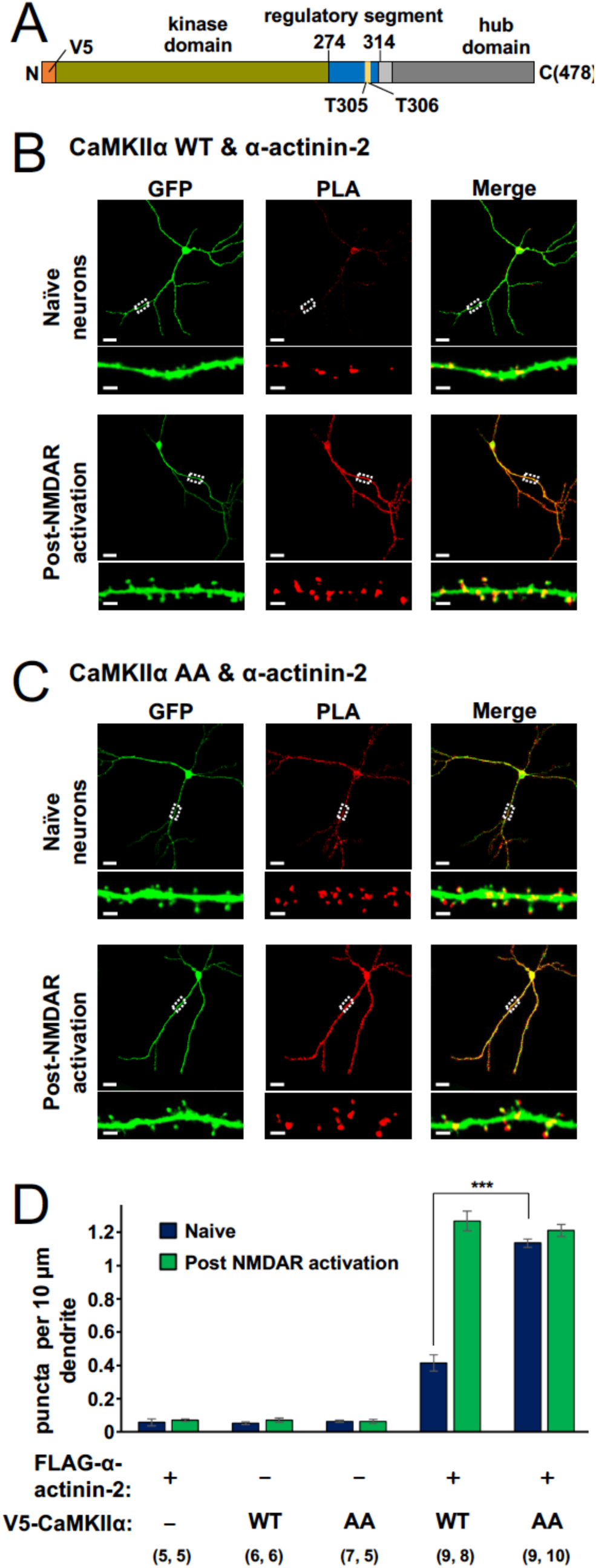
Basal interaction of CaMKII⍺ T305A/T306A with ⍺-actinin-2 is elevated. (A) Topology of V5-CaMKII⍺ highlighting location of putative regulatory threonines at positions 305/306. Panels (B) and (C) show anti-GFP immunofluorescence (left column) and anti-V5/anti-FLAG PLA puncta (middle column) in primary hippocampal neurons expressing GFP and FLAG-⍺-actinin-2 with either V5-CaMKII⍺ WT (B) or AA (C). In both cases imaging was performed either before (upper rows) or two hours after (lower rows) NMDAR activation. Scale bars are 20 𝜇m (square panels) and 2 𝜇m (dendrite close-ups). (D) Quantitation of anti-FLAG/anti-V5 PLA puncta per 10 𝜇m dendrite before (blue) and after (green) NMDAR activation in neurons expressing different CaMKII/actinin pairings. Data are presented as the mean ± standard error (SE). The number of neurons analysed for each condition is shown in parentheses with data collected from three independent cultures for all conditions. In panel D, data were analyzed using an unpaired two-tailed Student’s t-test (\*\*\**P* < 0.001).

A structural role for CaMKII was posited as soon as its neuronal abundance became apparent (19). CaMKII forms dodecamers (20, 21) with three domains per protomer capable of interacting with different proteins (22). Docking of CaMKII to NMDARs has emerged as a key initial step that occurs within the first ∼15 seconds of LTP induction (10). After Ca^2+^/CaM binds to the regulatory segment of CaMKII, a substrate motif in the GluN2B subunit tail is able to access the substrate-binding groove of the CaMKII kinase domain (23). The resulting complex is highly stable (24). The importance of this step was highlighted by a recent study that employed an ATP-competitive CaMKII inhibitor that is able to inhibit CaMKII without affecting its ability to bind NMDARs (4). Experiments with this compound show that, remarkably, CaMKII kinase activity is dispensable for LTP elicited by robust stimuli (4) – its structural capabilities are sufficient. ‘Follower’-type CaMKII interactions have been hypothesised as a means to translate the initial cue provided by CaMKII docking to NMDARs into the full structural and functional changes that occur in LTP (10). We recently discovered one such interaction that forms between CaMKII and α-actinin-2 within ∼1-2 minutes of the Ca^2+^ impulse (17). α-actinin-2 is an actin-crosslinking protein that is enriched in dendritic spines of excitatory synapses (25, 26). Either shRNA-mediated α-actinin-2 knockdown (27), or disruption of CaMKII-actinin interactions (17), prevents the formation of mushroom-shaped spines following NMDAR activation. α-actinin-2 is the only protein besides Ca^2+^/CaM known to bind to the CaMKIIα regulatory segment, but its access to the segment is partly occluded in inactive CaMKII (17). Association of CaMKII with GluN2B subunits releases the regulatory segment from the kinase domain, which enables higher affinity binding to the third and four EF hands (EF3-4) of α-actinin-2 (17). This likely explains why α-actinin-2 binding ‘follows’ initial docking to NMDARs (18).

As CaMKII residues T305 and T306 fall within the α-actinin-2 binding site (17, 28), we reasoned that the AA substitution might act at least in part by altering interactions between α-actinin-2 and CaMKII. In this study, we used *in situ* labelling to monitor interactions between α-actinin-2 and either wild-type (WT) or AA CaMKIIα in primary hippocampal neurons. We also analysed the effects of CaMKIIα AA expression on dendritic spine structure and synaptic currents. Calorimetry with purified fragments of the two proteins, and a crystal structure showing the structural basis of the interaction, explain how the AA substitution alters interactions between the two proteins. Our findings call for further revision of the molecular model for LTP, and further emphasise the structural rather than enzymatic function of CaMKII.

## Results

### Interactions between CaMKII**α** T305A/T306A and α-actinin-2 are elevated compared to those of WT CaMKII**α** in unstimulated neurons

We first aimed to determine whether association of CaMKIIα AA and α-actinin-2 is elevated *in situ* relative to wild-type (WT) kinase using proximity ligation assays (PLA). We have previously shown that PLA can be used to monitor interactions between CaMKII and actinin in primary hippocampal neurons, and that NMDAR activation triggers a marked increase in association of the two proteins (17). Rat primary hippocampal neurons were transfected on the tenth day *in vitro* (DIV10) with pIRES2-EGFP constructs for expression of FLAG-α-actinin-2 and V5-CaMKIIα variants to enable detection of actinin-CaMKII interactions by anti-FLAG/anti-V5 PLA after fixing on DIV14. Expression of either FLAG-α-actinin-2, V5-CaMKIIα WT or V5-CaMKIIα AA in isolation (**Fig. S1**) led to baseline levels of PLA puncta formation as expected. For neurons co-expressing FLAG-α-actinin-2 with V5-CaMKIIα WT, we detected 0.41 ± 0.05 PLA puncta per 10 μm dendrite (**Fig. 1***B*, upper row). For this WT pairing, puncta frequency rose to 1.27 ± 0.06 per 10 μm (*P* < 0.0001) following NMDAR activation with glycine (**Fig. 1***B*, lower low) – a chemical protocol that is considered a relatively realistic model of LTP (29, 30). These values are in line with our previous work (17). PLA puncta formation in naïve neurons co-expressing FLAG-α-actinin-2 and V5-CaMKIIα AA (**Fig. 1***C*, upper row) was strikingly elevated at 1.14 ± 0.05 puncta per 10 μm – 2.8-fold higher than the equivalent condition with WT CaMKII (*P* < 0.0001). However, for the AA pairing, NMDAR activation did not elevate puncta formation relative to naïve neurons (1.21 ± 0.04 puncta per 10 μm, **Fig. 1***C*, lower row). PLA puncta frequencies are summarized in **Figure 1***D*. Overall, our PLA imaging shows that interaction between CaMKIIα AA and α-actinin-2 is elevated in unstimulated neurons to a level only observed for WT kinase following the induction of chemical LTP.

### The T305A/T306A CaMKIIα substitution increases spine size without altering synaptic currents

Interactions between CaMKIIα and α-actinin-2 increase following NMDAR activation, and disruption of these interactions prevents formation of large ‘mushroom-type’ dendritic spines (17, 18, 27). Given that the AA variant of CaMKIIα associates with α-actinin-2 in the absence of NMDAR activation (**Fig. 1***C*, upper row), we hypothesised that spine morphology might be altered in naïve neurons expressing CaMKIIα AA. We determined the abundance of stubby (red), thin (amber), and mature mushroom-type (green) spines in primary hippocampal neurons expressing either FLAG-α-actinin-2 alone (**Fig. 2***A*, leftmost column), WT or AA V5-CaMKIIα variants alone (second-from-left and middle columns), or each CaMKIIα variant in combination with FLAG-α-actinin-2 (right-most two columns). We compared spine types in both naïve neurons (**Fig. 2***A*, upper row) and in neurons after induction of chemical LTP (lower row). Total spine densities were not statistically different between any of the five conditions before or after chemical LTP (∼3.5 spines per 10 μm, **Fig. S2**) but the abundance of particular spine types was influenced by expression of CaMKIIα AA (**Fig. 2***A* and *B*). Mushroom-type spines were in a minority in naïve neurons that expressed only WT CaMKIIα. Neurons expressing FLAG-α-actinin alone projected 0.58 ± 0.09 mushroom spines per 10 μm dendritic length; those expressing V5-CaMKIIα WT alone or in combination with actinin presented 0.55 ± 0.10 and 0.69 ± 0.06, respectively. Expression of V5-CaMKIIα AA alone triggered a 4-fold increase in mushroom spine abundance to 2.17 ± 0.18 per 10 μm (*P* < 0.001) compared to neurons expressing only V5-CaMKIIα WT. A similar effect was seen in neurons co-expressing V5-CaMKIIα AA and α-actinin-2, with mushroom spine abundance rising to 2.09 ± 0.19 per 10 μm (*P* < 0.001) relative to the equivalent WT condition (**Fig. 2***B*).

**Figure 2.**
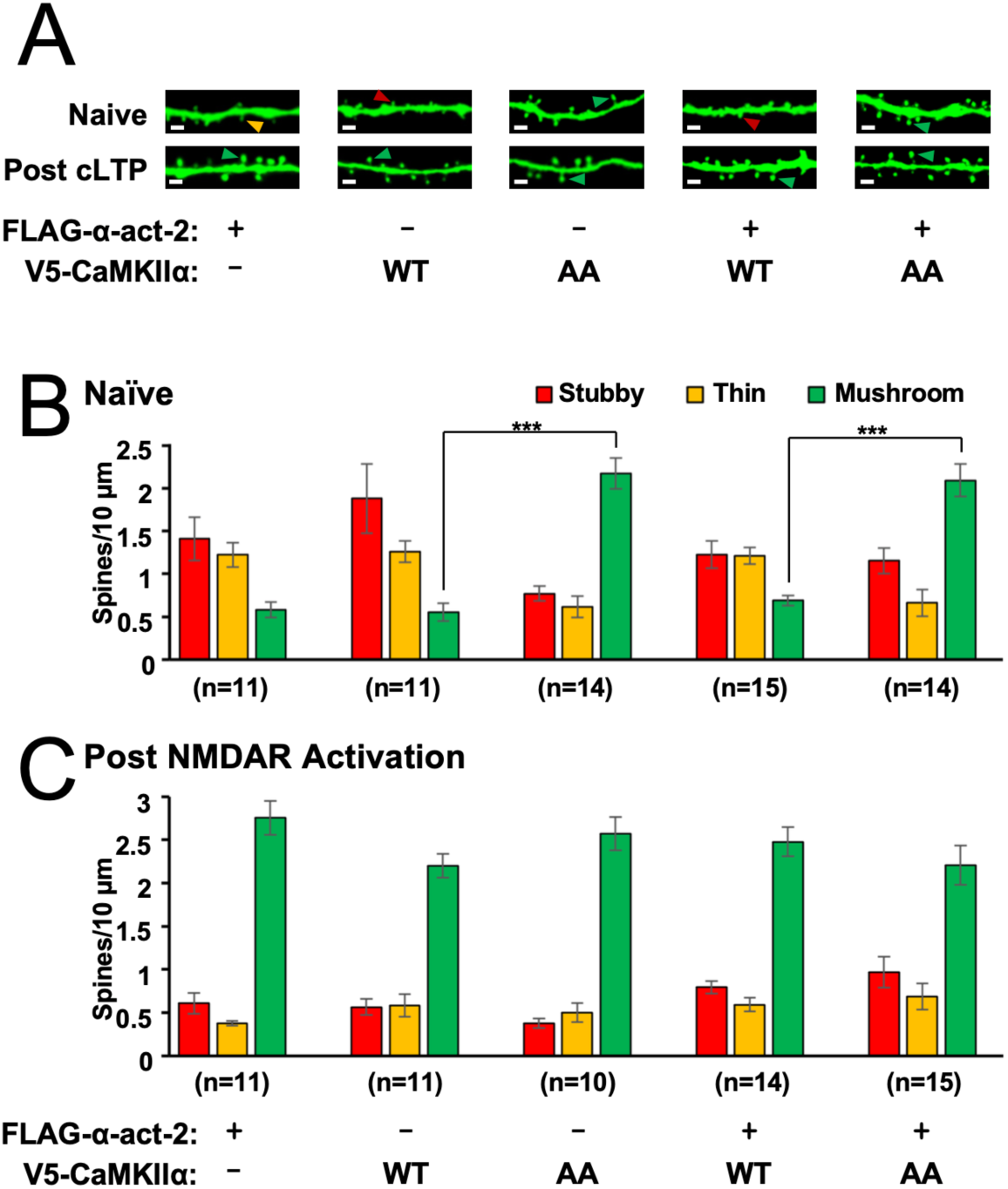
Effect of CaMKII⍺ T305A/T306A on spine morphology. (A) GFP imaging of dendrites in primary hippocampal neurons transfected with different combinations of pIRES2-GFP vectors expressing FLAG-⍺-actinin-2 or V5-CaMKII⍺ variants. Stubby (red), thin (orange), and mushroom (green) type spines are highlighted with arrows. Scale bars correspond to 2 𝜇m. Panels B and C show quantification of spine types across the three conditions either before (B) or after (C) NMDAR activation. Data are presented as mean ± SE spines per 10 𝜇m dendritic length. The number of neurons analysed for each condition is shown in parentheses. Neurons were imaged deriving from three independent cultures for each condition. Data were compared using unpaired two-tailed Student’s t-tests (\*\*\**P* < 0.001).

LTP is accompanied by an increase in spine width and proportion of spines with a mushroom-like structure, and this was evident in neurons expressing only WT CaMKIIα following NMDAR activation with both mushroom spine abundance (**Fig. 2***C*) and average spine width (**Fig. S2**) increasing as expected. After chemical LTP, mushroom spines had become predominant in neurons expressing FLAG-α-actinin-2 alone (2.76 ± 0.20 per 10 μm, **Fig. 2***C*), V5-CaMKIIα WT alone (2.20 ± 0.14), or V5-CaMKIIα WT in combination with α-actinin-2 (2.48 ± 0.17). Average spine width also increased for all three conditions: from 0.32 ± 0.01 to 0.57 ± 0.02 μm (α-actinin-2 only, *P* < 0.001), 0.33 ± 0.01 to 0.56 ± 0.01 (WT CaMKIIα only, *P* < 0.001), and 0.33 ± 0.01 to 0.56 ± 0.01 (WT CaMKIIα plus α-actinin-2, *P* < 0.001). However, spine morphology was little changed in neurons expressing CaMKIIα AA: neither spine type (**Fig. 2***C*) nor spine width (**Fig. S2***B*) was noticeably altered, with both of these characteristics now in line with the other experimental conditions. In sum, the spine imaging data indicates that the CaMKIIα AA substitution brings about morphological changes in dendritic spines resembling those that occur in neurons expressing WT kinase following LTP.

In wild-type neurons, the size of dendritic spine heads typically scales with the amplitude of miniature excitatory postsynaptic currents (mEPSCs) (31, 32). However, dissociation of these two attributes has been observed previously in neurons expressing mutated variants of CaMKIIα. For example, hippocampal primary neurons transfected with CaMKIIα T286D were found to develop larger spines but exhibited smaller EPSCs compared to WT controls (14). To determine whether the changes in spine architecture brought about by CaMKIIα AA were accompanied by changes in EPSCs, we recorded mEPSCs from untransfected neurons and neurons expressing either the WT or AA variant of CaMKIIα (**Fig. 3**). There was no difference in mean amplitudes for the three groups: mEPSCs were recorded in untransfected neurons with a mean amplitude of −22.0 ± 1.6 pA, (grey, **Fig. 3***A* and *B*) compared to −20.2 ± 1.6 pA for neurons transfected with WT CaMKIIα (orange), and −21.7 ± 1.6 pA for the AA variant (green, *P* = 0.93). mEPSC frequencies were variable within groups but again no differences were detected between the three groups (**Fig. 3***C* and **Table S1**). The same results were obtained when frequency and amplitude analysis was restricted to those (presumed proximal) mEPSCs having 10-90% risetimes of less than 1 ms (**Table S1**). Taken together, our data show that the CaMKIIα AA variant triggers increases in spine width and the proportion of mushroom-type spines without altering the mean amplitude of mEPSCs.

**Figure 3.**
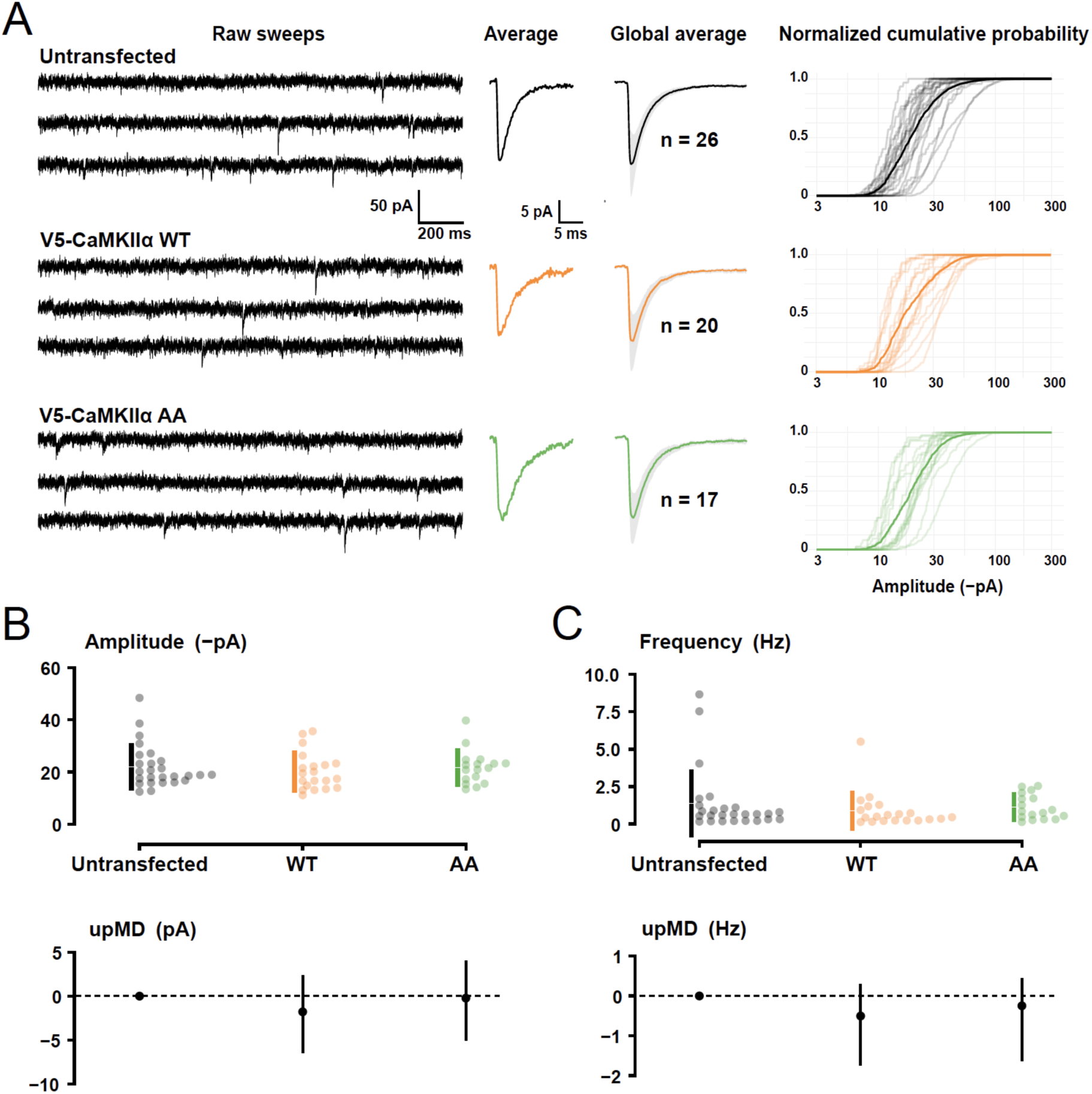
mEPSC recordings from primary hippocampal neurons. (A) Data from untransfected neurons (top row) or neurons transfected with either V5-CaMKIIα WT (middle row) or AA (bottom row). *Left*: Traces from three representative cells. *Middle*: the corresponding average mEPSC from each cell and the global average mEPSC (± SD, shaded) for each condition (untransfected, gray; V5-CaMKIIα WT, orange; V5-CaMKIIα AA, green). *Right*: Normalized cumulative probability plots of the amplitudes (log_10_ scale) for each group, with distributions from individual cells shown together with global averaged distributions (dark traces). The plots highlight the wide range within each group and the absence of marked differences between groups. (B) Cumming estimation plot for average mEPSC amplitudes (at −60 mV). The upper panel shows swarmplots with corresponding means and SDs indicated by gapped error bars. In the lower panel unpaired mean differences (upMD) from the untransfected condition are depicted as dots and each bootstrapped 95% confidence interval is indicated by the ends of the vertical error bars. (C) Same as B, but for mEPSC frequency.

### Calorimetry with purified proteins confirms that the T305A/T306A CaMKIIα variant binds tightly to α-actinin-2

Conventionally, effects of the AA substitution are attributed to decreases in phosphorylation at T305 and T306. In this case, one might attribute the increased association between α-actinin-2 and CaMKIIα brought about by the AA substitution to a decrease in phosphorylation at position T306 that occludes α-actinin-2 binding *in vitro* (28). However, previous studies have shown that one would not expect any resting phosphorylation at positions T286, T305 or T306 in unstimulated neurons (33). Furthermore, *in vitro* binding studies show that interactions between CaMKIIα and α-actinin-2 are not affected by phosphorylation at T305 (28), consistent with a crystal structure showing the interface between the two proteins (17). Therefore, one would expect any change in CaMKIIα – α-actinin-2 interaction resulting from suppression of inhibitory phosphorylation to be expressed in full by the single substitution T306A. However, single T306A substitution has no effect on *in situ* interactions between the two proteins (17). An alternative explanation is that the AA variant has an unexpected gain-of-function ability to bind tightly to α-actinin-2 irrespective of any differences in phosphorylation. To rigorously investigate this possibility, we performed isothermal titration calorimetry (ITC) with purified proteins. α-actinin-2 EF hands 3-4 (EF3-4, orange, **Fig. 4***A*) are the principal site for interactions with CaMKIIα (17, 28). We previously showed that EF3-4 binds isolated WT CaMKIIα regulatory segment (294–315) with K_d_ = 32 ± 1 µM, whereas interactions between EF3-4 and a CaMKIIα construct that includes both the kinase domain and regulatory segment (1–315) are so weak as to be undetectable by ITC. Here, we performed equivalent measurements using CaMKIIα fragments bearing single or double alanine substitutions at T305 and T306. Measurements with single alanine substitutions yielded similar results to the WT sequence. EF3-4 bound to T305A regulatory segment with K_d_ = 19.4 ± 0.7 μM, and to the T306A equivalent with K_d_ = 19.8 ± 1.9 μM (**Fig. S3**). Furthermore, it was not possible to detect interactions between EF3-4 and CaMKIIα T305A or T306A in the context of the longer 1-315 construct (**Fig. 4***B* and *C*), recapitulating the result with WT construct (17). Remarkably, CaMKIIα (1–315) AA bound tightly to EF3-4 with K_d_ = 154 ± 9 nM (**Fig. 4***D*), which is 100-fold tighter than even the interaction between EF3-4 and isolated WT regulatory segment. Surprisingly, though, the AA substitution did not markedly alter interactions in the context of the isolated regulatory segment with interaction to EF3-4 occurring with K_d_ = 14.7 ± 1.2 μM in this case (**Fig. S3**). This suggests that high affinity interactions between the AA variant and α-actinin-2 involve CaMKIIα elements beyond positions 294-315. Full thermodynamic parameters obtained for all ITC measurements are shown in **Table S2**. Overall, the ITC data reveal that inactive CaMKIIα bearing the AA substitution can bind α-actinin-2 with remarkably high affinity, and that alanine substitutions at both T305 and T306 are necessary to bring about this effect.

**Figure 4.**
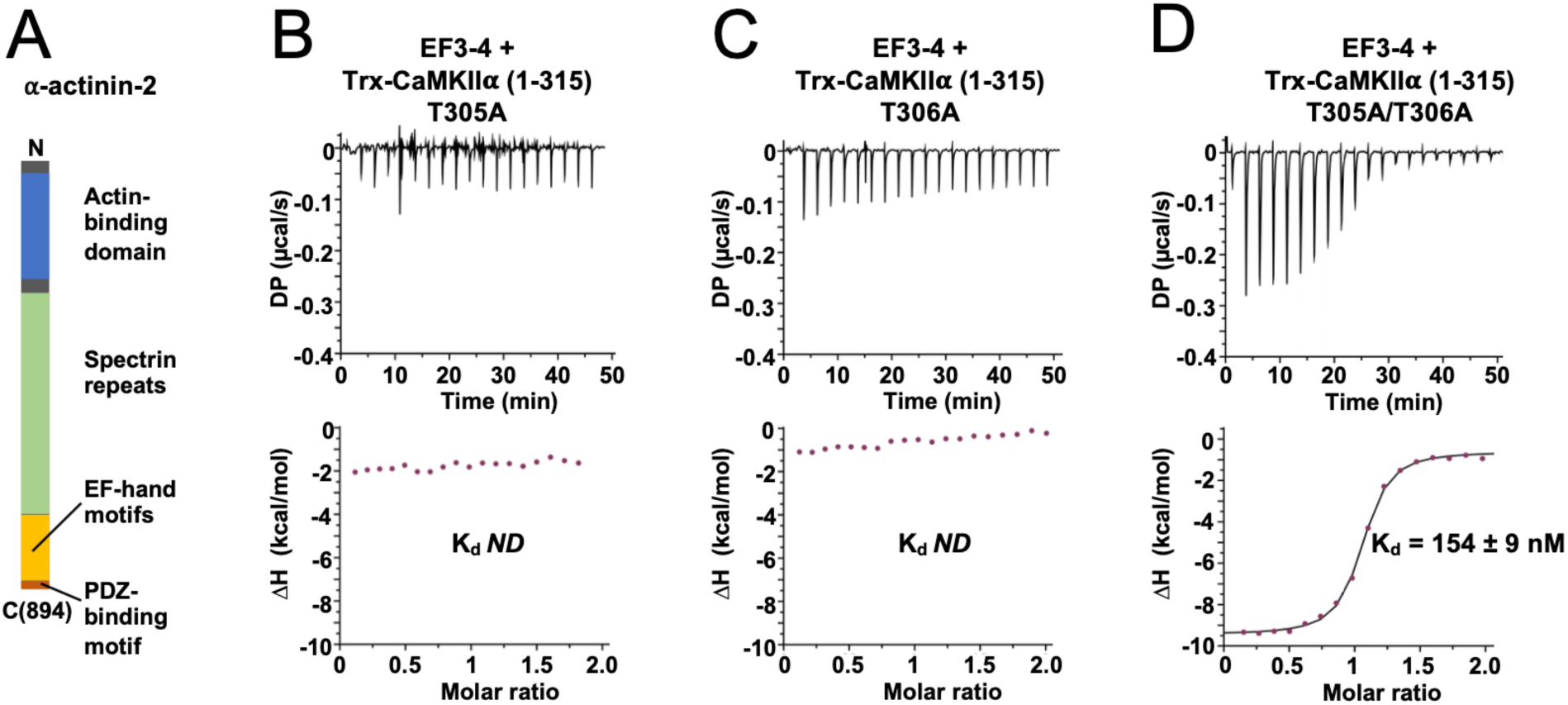
⍺-actinin-2 binds tightly to a CaMKII⍺ AA construct spanning residues 1-315. (A) Topology of ⍺-actinin-2 with the EF3-4 region highlighted in orange. Panels B-D show representative isotherms for binding of ⍺-actinin-2 EF3-4 to Trx-CaMKII⍺ (1-315) constructs bearing either T305A (B), T306A (B), or T305A/T306A (D) substitutions. The top sub-panels show the raw power outputs over time (µcal/s); the bottom sub-panels show the integrated heat changes including a line of best fit to a single site binding model. Stated K_d_ values are mean ± SE from experimental replicates. ND = not determined.

### Molecular basis of high affinity association between α-actinin-2 and the T305A/T306A variant

CaMKIIα (1–315) AA bound to EF3-4 tightly, whereas a peptide spanning residues 294-315 of CaMKIIα AA did not. This indicates that the AA variant binds α-actinin-2 using an unanticipated binding mode. To understand the molecular basis of the interaction, we determined crystal structures of a complex between EF3-4 and CaMKIIα (1–315) AA (**Fig. 5***A*). Structures were determined with either no nucleotide (PDB ID 7B55; 1.6 Å resolution, **Fig. S4***A*), with Mg^2+^/AMP-PNP (PDB ID 7B56; 1.45 Å resolution, **Fig. S4***B*), or with Mg^2+^/ADP (PDB ID 7B57; 1.95 Å resolution, **Fig. S4***C*). Full crystallographic statistics are shown in **Table S3**. In the nucleotide-free complex, a MES molecule occupies the nucleotide-binding site (**Fig. S4***A*). The three structures are broadly similar (**Fig. S4***D*), with RMSD for all Cα positions of 0.63 Å (7B55-7B56), 0.53 Å (7B55-7B57), and 0.371 Å (7B56-7B57). The structures are most different in the vicinity of the nucleotide-binding pocket (**Fig. S4***E-G*), but α-actinin-2 and CaMKIIα AA interact in the same way in all three cases. Our analysis focuses on the complex with ADP (7B57). In the complex, CaMKIIα kinase domain (green, **Fig. 5***B*) adopts a similar conformation to that observed in other structures of auto-inhibited CaMKIIα (20). For example, alignment of CaMKIIα positions 7-300 between the Mg^2+^/ADP complex structure solved here and the equivalent region in a full-length autoinhibited CaMKIIα construct (3SOA) (20) gives RMSD of 0.79 Å. Electron density is visible for the CaMKIIα regulatory segment up to residue M307, with the regulatory segment exiting the kinase domain in the vicinity of the glycine rich nucleotide-coordinating loop (**Fig. 5***B*). The α-actinin-2 EF3-4 region (orange, **Fig. 5***B*) forms an extensive interface with the CaMKIIα regulatory segment (blue), which presents a series of aliphatic sidechains on the side opposite to the kinase domain including A295, L299, A302, and I303 (**Fig. 5***C* and *D*). Van der Waals interactions between these amino acids and EF3-4 residues F835, V831, L854, and C862 make up the core of the interface (**Fig. 5***C* and *D*). A further hydrophobic interaction occurs between the sidechains of M307_CaMKII_ and Y861_EF3-4_ (**Fig. 5***D*). The interface is supported by hydrogen bonds at either end: the amine group of K292_CaMKII_ H-bonds with the main-chain oxygen of I837_EF3-4_ (**Fig. 5***C*); and the main-chain oxygen of A302_CaMKII_ H-bonds with the hydroxyl group of Y889_EF3-4_ (**Fig. 5***C* and *D*). Finally, the interface includes a salt-bridge between the sidechains of R296_CaMKII_ and E853_EF3-4_ (**Fig. 5***D*).

**Figure 5.**
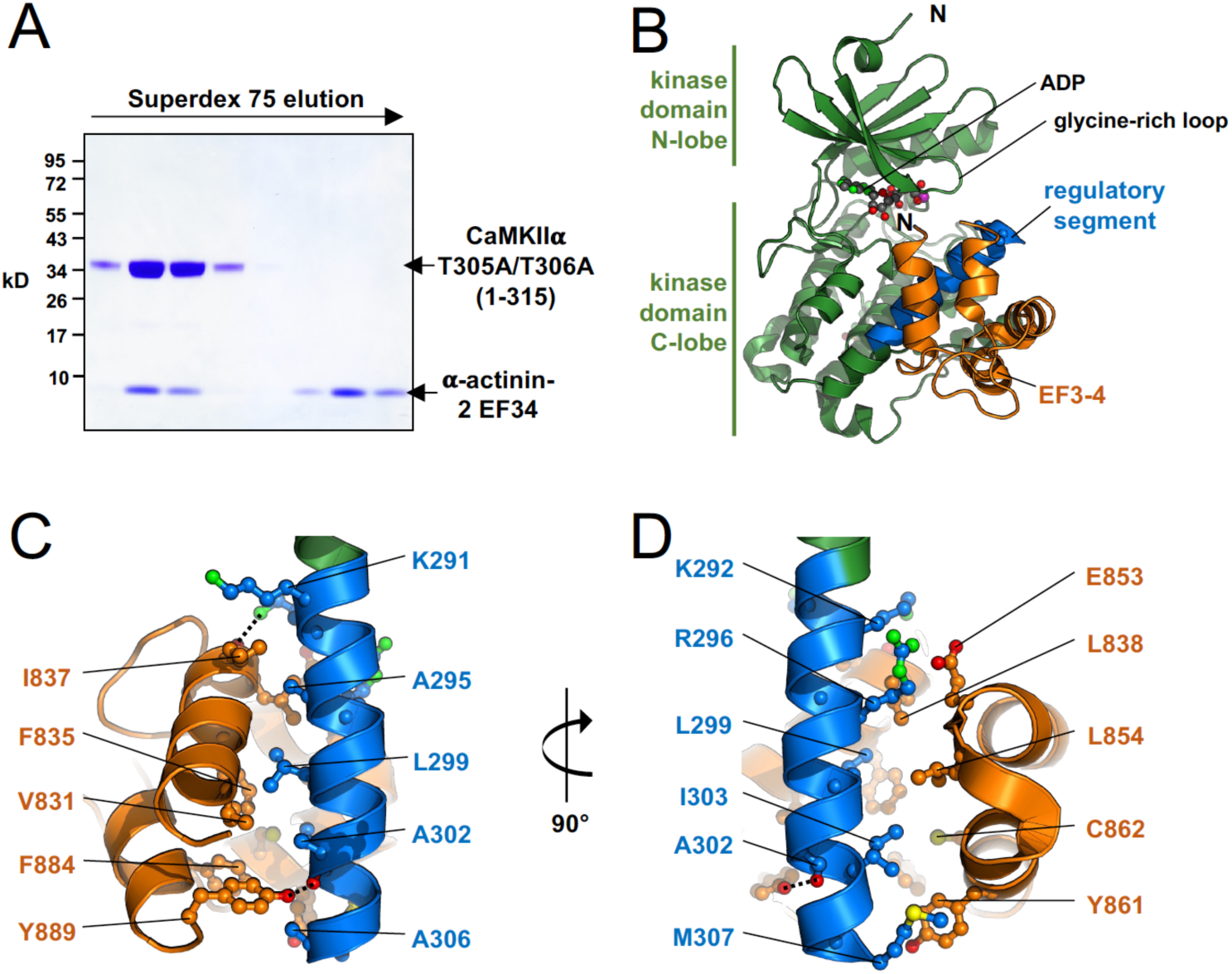
Crystal structure of complex between ⍺-actinin-2 EF3-4 and CaMKII⍺ AA. (A) Coomassie gel showing elution of a mixture of CaMKII⍺ 1-315 AA and a molar excess of ⍺-actinin-2 EF3-4 from a Superdex 75 size exclusion column. The earlier fractions containing the complex were used for crystallization. (B) Overview of the structure obtained following crystallization with Mg^2+^/ADP (PDB ID 7B57). The kinase domain and regulatory segment of CaMKII⍺ are shown in green and blue, respectively; EF3-4 is colored orange. Panels C and D show two close-up views of the core interface related by a 90° rotation, with ⍺-actinin-2 residues in orange and CaMKII residues in blue.

Comparison to structures of WT CaMKIIα regulatory segments in complex with α-actinin-2 and CaM reveals that the AA substitution supports a binding mode that has not been observed before. **Fig. 6** shows aligned structures of WT CaMKII regulatory segment in complex with Ca^2+^/CaM (PDB 2WEL, **Fig. 6***A*) (34) and α-actinin-2 EF3-4 (PDB 6TS3, **Fig. 6***B*) (17) alongside the structure of AA-variant regulatory segment in complex with EF3-4 (**Fig. 6***C*) extracted from the Mg^2+^/ADP structure solved in this study (PDB 7B57). The CaM complex involves the δ isoform of CaMKII (**Fig. 6***B*) but this isoform is identical in the regulatory region corresponding to positions 282-315 in CaMKIIα. Equivalent side-on views of the CaMKII regulatory segment are shown in **Fig. 6***A-C*, with each structure centred on T305. A perpendicular view along the axis of the regulatory segment α-helix is also shown in each case (**Fig. 6***A-C*, right-hand sub-panels). Ca^2+^/CaM fully envelopes the CaMKII regulatory segment (**Fig. 6***A*), which explains why CaM binds much more tightly to isolated regulatory segment or pT286-activated CaMKII where steric hindrance by the kinase domain is absent or reduced (35, 36). EF3-4 binds predominantly to the C-terminal part of WT CaMKIIα regulatory segment (**Fig. 6***B*), with direct interactions to CaMKII spanning positions L299 to F313. EF3-4 is rotated by ∼50° relative to the third and fourth EF hands of CaM around the axis of the regulatory segment helix, with the third EF hand positioned in the foreground according to the view in **Fig. 6** (right-hand column) in both cases. Structural alignment of isolated CaMKII regulatory segments or EF3-4 domains from the WT and AA complexes shows that each separate polypeptide retains a highly similar conformation in both complexes (**Fig. S5**). However, the relative orientation of the two binding elements is markedly altered (**Fig. 6***B* and *C*). In the complex with CaMKIIα (1–315) AA, EF3-4 binding is shifted towards the N-terminus of the regulatory segment (**Fig. 6***C*) with direct interactions extending up to CaMKII position K291 and only reaching M307 in the C-terminal direction. Most strikingly, EF3-4 is flipped according to the view in **Fig. 6***A-C* (right-hand column) with the fourth EF hand (light orange) in the foreground unlike the other two complexes.

**Figure 6.**
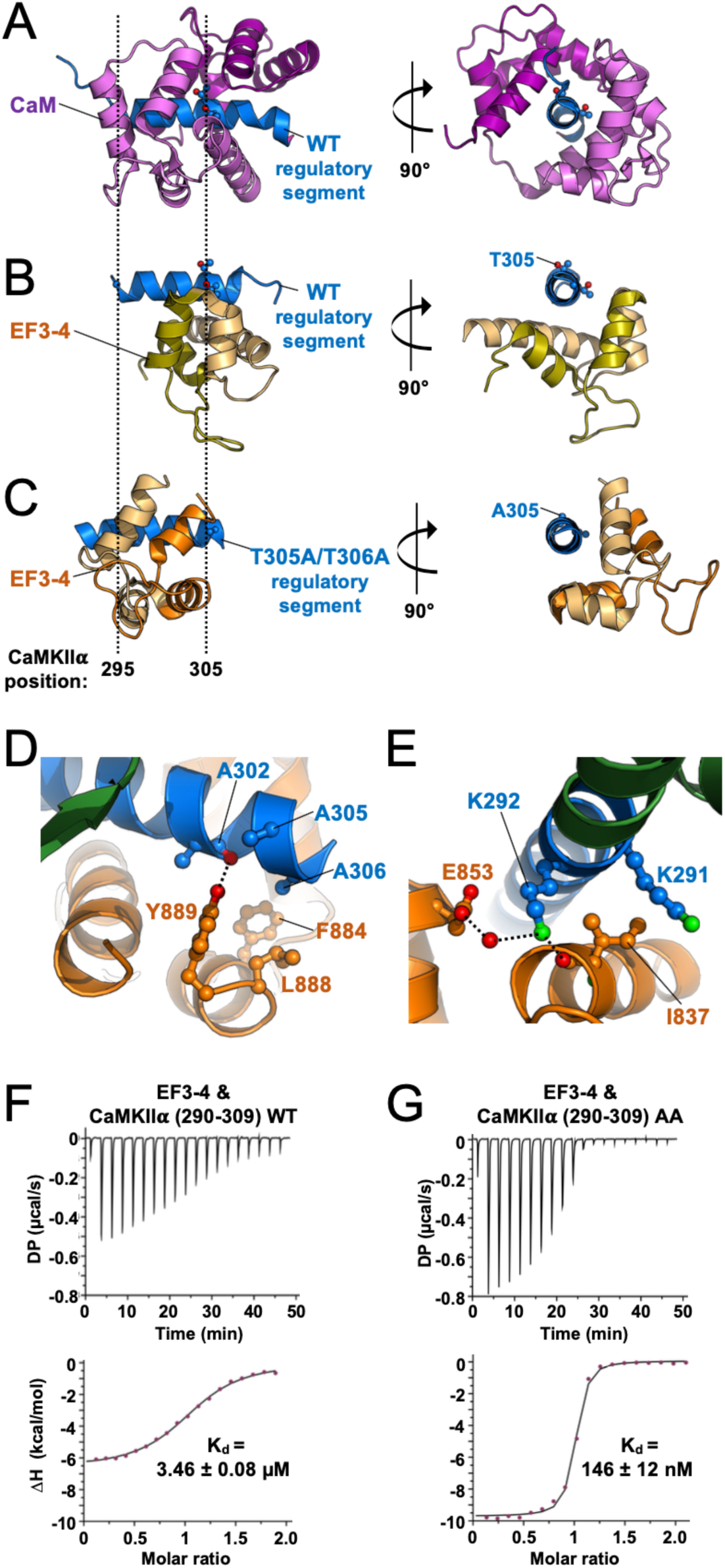
Molecular basis of enhanced ⍺-actinin-2 binding to CaMKII⍺ AA. Panels A to C show equivalent views of the CaMKII⍺ regulatory segment (blue) highlighting the relative binding orientations of CaM (A) and EF3-4 (B) to WT regulatory segment, and EF3-4 to the AA variant (C). The three panels correspond to structures 2WEL, 6TS3 and 7B57, respectively. The CaM N-lobe is coloured purple: the C-lobe is violet. For ⍺-actinin-2, EF3 is coloured light orange; EF4 is olive. Panels D and E show close-up views of the the EF3-4 – CaMKII⍺ AA interface highlighting interactions in the vicinity of A305/A306 (D), and interactions involving K291/K292 (E). Panels F and G show representative isotherms for binding of ⍺–actinin-2 EF3-4 to either the WT (F) or AA (G) variants of CaMKII⍺ 290-309. The top sub-panels show the raw power output (µcal/s) over time; the bottom sub-panels show the integrated data including a line of best fit to a single site binding model. Stated K_d_ values are averages from three replicates.

The much higher affinity of EF3-4 for CaMKIIα AA residues 1-315 compared to 294-315 (**Figs. 4***D* and **S3***C*) led us to suspect that EF3-4 contacts elements within the kinase domain of the AA variant. However, the structure reveals no direct interactions involving the kinase domain (**Fig. S5***C*). Instead, substitution of alanine at positions 305 and 306 enables EF3-4 to invert its third and fourth EF hands and mediate a different network of interactions to the regulatory segment with a shift of approximately two helical turns towards the N-terminus of the segment. In this new binding mode, A306 mediates van der Waals interactions with L888_EF3-4_ and F884 _EF3-4_ and the substitution at position 305 provides access for a H-bond between the hydroxyl group of Y889_EF3-4_ and the main-chain carbonyl of A302_CaMKII_ (**Fig. 6***D*). At the N-terminal end of the segment, the side-chain of K291_CaMKII_ packs against I837_EF3-4_ (**Fig. 6***E*). In addition, the terminal amine of K292_CaMKII_ H-bonds with the main-chain oxygen of I837_EF3-4_ and also forms a water-mediated H-bond with the side-chain of E853_EF3-4_. To confirm that the higher affinity of α-actinin-2 for the AA CaMKIIα variant results from these additional interactions, we performed further ITC measurements with N-terminally shifted regulatory segment peptides (**Fig. 6***F* and *G*). Consistent with the binding mode observed in the structure, EF3-4 bound to AA CaMKIIα 290-309 more than 20-fold more tightly (K_d_ = 146 ± 12 nM, **Fig. 6***G*) than to the WT equivalent (3.46±0.08 µM, **Fig. 6***F*) with a dissociation constant comparable to the 1-315 construct (**Fig. 4***D*, Kd = 154 ± 9 nM). In sum, our structural and ITC data show that the AA substitution allows α-actinin-2 to adopt a different higher-affinity binding mode that involves extensive interactions with N-terminal elements of the CaMKII regulatory segment.

## Discussion

In this study, we show that the AA variant of CaMKIIα has a gain-of-function ability to bind tightly to α-actinin-2 (**Fig. 1***C*); based on ITC measurements the binding affinity between an inactive CaMKIIα construct and the EF3-4 region of α-actinin-2 is ∼150 nM (**Fig. 4***D*). The EF3-4 region of α-actinin-2 adopts a different binding mode when associating with the AA variant, and, unlike the WT sequence (17), its interface on CaMKIIα is fully accessible when the kinase is inactive (**Fig. 5**). Expression of the AA variant in cultured neurons increases the proportion of mushroom-type dendritic spines in unstimulated neurons (**Fig. 2***B*). Previous studies have shown that α-actinin-2 is essential for the development of enlarged mushroom-type spines during LTP through interactions with CaMKIIα (17, 27). Our findings suggest that the AA variant provides a shortcut to this structural component of LTP by interacting directly with α-actinin-2, whereas in WT neurons the interaction only forms after NMDAR activation (17).

Our findings are consistent with a study that employed the CaMKIIα AA variant for purified α-actinin-2 pull-downs and found these to be more robust than for the WT kinase (28). Prior work indicates that phosphorylation of T305 but not T306 can potentially inhibit interactions between CaMKII and α-actinin-2 (28). T305 projects away from the interface between WT CaMKII and α-actinin-2 (17), and prolonged *in vitro* incubation of autonomously-active T305A (but not T306A) CaMKII reduces α-actinin-2 pull-down with CaMKIIα (28). On this basis, one would expect any increase in CaMKII-actinin association deriving from suppression of inhibitory phosphorylation to be expressed in full by the T305A variant. However, we found previously that T305A CaMKIIα behaved like WT kinase, with α-actinin-2 interactions increasing markedly after NMDAR activation (17). Therefore, the most logical explanation is that the effects mediated by the AA variant derive from its ability to bind tightly to α-actinin-2 rather than through suppression of inhibitory phosphorylation. Unanticipated effects of alanine substitutions have been documented previously. For example, the active site mutant C146A form of Ubp8 unexpectedly binds tightly to, and sequesters, ubiquitin (37). Synergistic effects of multiple alanine substitutions have been observed in a screen of alanine-substituted peptide binders to the oncogenic protein MDM2 (38), and alanine substitutions at three positions in an inhibitor peptide derived from CaMKIIN also unexpectedly increased potency (39).

The gain-of-function ability to bind tightly to α-actinin-2 provides an alternative explanation for the effects of the AA substitution in CaMKIIα but does not rule out the possibility that phosphorylation at T305 and/or T306 is important. What then is the evidence that these sites are phosphorylated? CaMKII preferentially phosphorylates serine/threonine residues when arginine is present 3 residues prior in the primary sequence (the ‘-3’ position): mutation of the −3 arginine in a model substrate peptide led to a ∼300-fold decrease in catalytic efficiency including an ∼80-fold decrease in V_max_ (40). Furthermore, the presence of a non-hydrophobic residue at the −5 position greatly decreases phosphorylation (40). According to these criteria, one would expect T286 to be a good substrate (<u>M</u>H<u>R</u>QE**T**), T305 to be an extremely poor substrate (<u>K</u>G<u>A</u>IL**T**), and T306 to be a poor substrate (K<u>G</u>AILT**T)**. A quantitative proteomic study performed with CaMKIIα purified from insect cells fits with what would be expected from the primary sequence: Baucum and co-workers measured phosphorylation at specific sites before and after Ca^2+^/CaM stimulation, and also in a third phase dependent on autonomous activity after Ca^2+^/CaM removal (16). T286 was robustly phosphorylated in the second phase (∼80 % sites phosphorylated), whereas T306 phosphorylation was elevated to ∼10 % after the third phase and T305 phosphorylation could not be detected under any circumstances (16). Interestingly, phosphorylation at S314 was elevated to ∼ 40 % for S314 after the third phase – this position was actually considered in early studies investigating inhibitory CaMKII phosphorylation (11). For inhibitory phosphorylation at T305 or T306 to meaningfully inhibit subsequent Ca^2+^/CaM activation, one would expect that it would occur at higher levels than observed in this study under idealised circumstances. An alternative role for T305/T306 phosphorylation is the targeting of CaMKII to inhibitory synapses (33) – this mechanism is more compatible with inefficient T305/T306 phosphorylation since phosphorylation of, e.g., T306 in a single CaMKII protomer could be sufficient for targeting of a full dodecamer. In general, it is difficult to reliably monitor phosphorylation at specific sites in cells. Many studies of T305/T306 phosphorylation have relied on antibodies raised against short phosphopeptides. However, typically such antibodies are not validated using lysates from knock-in animals bearing substitutions at the target residues. Furthermore, while phospho-specific antibodies can reveal increases in phosphorylation relative to baseline they do not reveal what proportion of a given site has been phosphorylated. Interestingly, a study employing Phos-tag band shifting to monitor AMPAR phosphorylation in neuronal extracts suggests that less than 1 % GluA1 subunits are phosphorylated even after chemical LTP (41). This calls into question the significance of phosphorylation of these subunits at position S831 and S845 (10). Mass spectrometry (MS)-based quantitation of the absolute abundance of specific proteins within the postsynaptic density was essential for establishing a realistic model of this synaptic sub-structure (42, 43). It is much more challenging to apply proteomics to quantify phosphorylation levels at specific amino acids, but studies of this type (44) would help to distinguish which specific sites are important during synaptic plasticity.

CaMKIIα T305A/T306A substitutions are frequently combined with the phospho-mimetic change T286D – this ‘D/AA’ variant is considered as a ‘constitutively-active’ form of CaMKII that both increases the size of dendritic spines and increases AMPAR current amplitudes with no need for a Ca^2+^ impulse (9, 13, 14, 45–50). While CaMKIIα D/AA increases AMPAR current amplitudes, expression of the T286D substitution alone has the opposite effect (14). One interpretation of this result is that when threonine is present at positions 305 and 306, CaMKIIα auto-phosphorylates at these sites due to its autonomous activity, which prevents its full activation by Ca^2+^/CaM (14). However, this explanation doesn’t account for the finding that the T286D substitution also depresses AMPAR currents when introduced in combination with the kinase-inactivating mutation K42R (14). Experiments with Camui-α, which reports on the activation state of CaMKIIα, also cast doubt on the conventional interpretation (9). The basal fluorescent lifetime of Camui-α was found to be elevated in variants that included the T286D substitution either alone or in combination with the AA substitution (9). However, both forms showed a small but similar increase in activation upon glutamate uncaging indicating that the AA substitution does not noticeably alter the ability of Ca^2+^/CaM to activate CaMKII bearing the T286D mutation. Our finding that the AA substitution binds tightly to α-actinin-2 provides a more congruent explanation for the strongly potentiating behaviour of the D/AA mutation. α-actinin-2 is enriched in dendritic spines, and it interacts with several core postsynaptic proteins including NMDARs (51), PSD-95 (52), and densin-180 (53). Targeting autonomously-active CaMKII to this compartment may accurately mimic LTP, in which stable interactions between CaMKII and proteins including NMDARs and actinin are a key feature (4, 18). Interestingly, we found that CaMKIIα AA expression did not alter the amplitude of mEPSCs in cultured hippocampal primary neurons (**Fig. 3***D* and *E*). This suggests that the actinin-CaMKII interaction can mediate structural changes – likely via remodelling of the actin cytoskeleton (18) – without increasing AMPAR currents, and implies that interactions involving the CaMKII kinase domain are required for the full expression of LTP.

The importance of structural rather than enzymatic protein function is an emerging theme in the mechanics of synaptic plasticity (10). Recent work shows that CaMKII docking to NMDARs is essential for LTP whereas LTP proceeds normally if an activity-selective inhibitor is applied shortly after a strong Ca^2+^ impulse (4). Experiments with a different abundant PSD protein reinforce this theme: mutations within the GTPase-activating protein (GAP) domain of SynGAP do not inhibit synaptic plasticity (54). Instead, SynGAP is thought to modulate synaptic strength by physically competing with AMPAR-TARP complexes for recruitment to the core PSD vertical filament protein PSD-95 (54). The ability of CaMKII to render itself autonomously active via auto-phosphorylation within dodecamers is a remarkable and unique characteristic that was reasonably considered as a mechanism for long-term memory storage (55). However, imaging with reporters of CaMKII conformation shows that kinase activity only lasts for a matter of seconds after NMDAR activation (56, 57). Consistent with these studies, CaMKII photo-inhibition has no effect on LTP if delayed until one minute after LTP induction (58). The function of T286 phosphorylation is now thought to be limited to the initiation phase including (59) detecting specific Ca^2+^ signal frequencies in LTP induction while playing no role in LTP maintenance (9). Our findings further downplay phosphorylation-based regulation of CaMKII activity and reinforce the concept that highly abundant synaptic enzymes can operate primarily through their ability to mediate protein-protein interactions.

## Materials and Methods

### Protein expression and purification

Human α-actinin-2 EF3-4 (positions 827-894) was expressed with an N-terminal Tev-cleavable His-GST tag within pET28. Protein expression was induced in Rosetta plysS *E. coli* with 1 mM IPTG, and cells were harvested after overnight incubation at 20 °C. Cells were lysed in Ni-NTA buffer A (500 mM NaCl, 25 mM Tris pH 8, 1 mM Benzamidine, 30 mM imidazole) supplemented with 0.1 mg/mL lysozyme and one cOmplete protease inhibitor tablet per 100 mL, then clarified by high-speed centrifugation following sonication. His-GST-EF3-4 was enriched by sequential Ni-NTA agarose (Qiagen) and Glutathione Sepharose 4B (Cytiva) affinity capture as before (17) prior to overnight cleavage with Tev protease at 4 °C. Finally, EF3-4 was resolved from cleaved GST using a HiLoad 16/600 Superdex 75 column equilibrated in 20 mM HEPES, pH 7.5, 0.15 M NaCl, and 1 mM DTT. CaMKIIα 1-315 variants were expressed with Tev-cleavable N-terminal His-Trx tag in Rosetta (DE3) pLysS *E. coli* (Merck) using pNH-TrxT vector as before (17). Alanine substitutions at positions T305 and T306 were introduced by site directed mutagenesis using primers listed in **Table S4**. Trx-CaMKII construct expression was induced with 0.2 mM IPTG, with cells harvested after overnight incubation at 18 °C. Initial purification was performed by affinity to Ni-NTA agarose and anion exchange with Q Fast Flow columns (Cytiva) as before (17). We found that if the Trx moiety was removed by Tev cleavage, CaMKIIα 1-315 was prone to precipitate unless bound to EF3-4. Therefore, different purification strategies were adopted for crystallography and ITC. For ITC measurements, the Trx tag was retained, and Trx-CaMKIIα was immediately subjected to size exclusion with a HiLoad Superdex 75 column (Cytiva) equilibrated in gel filtration buffer (20 mM HEPES pH 7.5, 150 mM NaCl, 1 mM DTT) following anion exchange. For crystallisation of the complex, the His-Trx-CaMKIIα AA 1-315 construct was mixed with a two-fold molar excess of EF3-4 for one hour, before addition of Tev protease. The complex was then resolved from excess EF3-4 and Trx using the same size exclusion approach on the following day. All protein samples were concentrated using 10K MWCO centrifugal concentrators (Sartorius). Dialysis was performed using Slide-a-Lyzer 2K MWCO cassettes (Thermo Scientific).

### Crystallography

For crystallisation of CaMKIIα 1-315 AA in complex with α-actinin-2 EF3-4, the CaMKII construct was first mixed with a ∼2-fold molar excess of EF3-4 before the 1:1 complex was separated from excess EF3-4 using size exclusion with a Superdex 75 column equilibrated in 20 mM HEPES, pH 7.5, 0.15 M NaCl, 1 mM DTT. The complex was concentrated to 10 mg/mL and crystals were grown using sitting drop vapor diffusion with precipitant solution containing 0.2 M Lithium sulfate, 0.1 M MES pH 6.0, 20 % w/v PEG 4000. For crystals containing ADP or AMP-PNP, the precipitant solution was supplemented with 10 mM MgCl_2_ and 5 mM nucleotide. Diffraction data were collected at Diamond Light Source beamline I24 and reduced using DIALS (60), before scaling with Aimless (61). The structures were solved by molecular replacement using Phaser (62), and finally refined in PHENIX (63). Full data collection and refinement statistics are provided in **Table S3**. Structural alignments and RMSD calculations were performed using GESAMT (64).

### Isothermal titration calorimetry

All ITC measurements were collected with a MicroCal PEAQTM (Malvern Panalyticial). Interactions with CaMKIIα peptides were performed at 25 °C in 25 mM HEPES pH 7.5 and 150 mM NaCl, with injections from a syringe containing 500 μM peptide into a cell containing 50 μM EF3-4. For interactions between Trx-CaMKIIα 1–315 variants and EF3-4, 2 mM MgCl_2_ and 1 mM ADP were added to the buffer, and in this case the syringe was filled with 300 μM EF3-4 while the cell contained 30 μM Trx-CaMKIIα 1–315. In all cases, data was collected with 2 μL injections at 2 min intervals and constant mixing at 750 rpm. Bicinchoninic acid assays and A280 absorbance were used in tandem to determine protein concentrations. The same EF3-4 preparations were used for the ITC measurements presented in this study and previous measurements with WT CaMKII (17). ITC data was collected and processed using MicroCal Origin software (Malvern Panalytical), using non-linear least-squares fitting to single binding models to estimate thermodynamic parameters.

### Hippocampal neuron culture and chemical LTP

Each hippocampal culture was a mixed population derived from single litters of ∼5 E18 Sprague Dawley pups. Dissociated hippocampal neurons were plated on 13 mm glass coverslips that had been pre-treated with poly-L-lysine (1 mg/mL) at 1×10^5^ cells per coverslip. Neuronal cultures were maintained in neurobasal medium supplemented with B27, GlutaMAX, and Penicillin/Streptomycin. Neurons were transfected on DIV10 using 0.8 μg DNA and 2 μL Lipofectamine-2000 per coverslip. N-FLAG-α-actinin-2 and N-V5-CaMKIIα variants were both expressed using pIRES2-GFP vectors. The AA variant was generated using site-directed mutagenesis with primers T305A&T306A_F and R (**Table S4**). For double transfections, 0.4 μg of each vector was included in the mixture. Cultures were maintained until chemical LTP and fixing on DIV14. Chemical LTP (cLTP) was induced by activating NMDARs with glycine (30, 65). Neurons were first transferred into control solution (5 mM HEPES pH 7.4, 125 mM NaCl, 2.5 mM KCl, 1 mM MgCl_2_, 2 mM CaCl_2_, 33 mM D-glucose, 20 μM D-AP5, 3 μM strychnine, 20 μM bicuculline, 0.5 μM TTX) for 20 min at room temperature before cLTP was induced by 10-min incubation in cLTP solution (5 mM HEPES pH 7.4, 125 mM NaCl, 2.5 mM KCl, 2 mM CaCl_2_, 33 mM D-glucose, 3 μM strychnine, 20 μM bicuculline, 0.2 mM glycine). Subsequently, neurons were returned to control solution to allow structural changes to develop before fixing in PBS supplemented with 4% paraformaldehyde, 4% sucrose, and 0.2% glutaraldehyde. Neurons were fixed two hours after induction of cLTP for PLA imaging, and four hours after cLTP induction for spine width analysis. Experiments involving rats were performed in accordance with the United Kingdom Animals Act, 1986 and within University College London Animal Research guidelines overseen by the UCL Animal Welfare and Ethical Review Body under project code 14058.

### Confocal imaging and PLA

For spine width analysis, fixed neurons on glass coverslips were permeabilised for 5 min at RT in PBS supplemented with 1 % BSA/0.1 % Triton X-100, blocked for 1 hour in PBS supplemented with 10 % BSA and incubated overnight at 4 °C in 1 % BSA-PBS containing chicken anti-GFP (RRID_Ab: 300798, 1:500 dilution). On the following morning, neurons were washed and incubated for a further hour in 1 % BSA-PBS containing rabbit anti-chicken Alexa Fluor 488 (RRID_Ab:2339327. 1:500 dilution). After washing in PBS, coverslips were mounted on glass slides with ProLong Gold antifade mountant (Thermo Fisher) and sealed using nail varnish. PLAs were performed using reagents from a Duolink In Situ PLA kit. For PLAs, fixed neurons were permeabilised and blocked in the same way prior to overnight incubation with 1 % PBS containing the following primary antibodies: mouse anti-V5 (RRID_Ab: 10977225, 1:500 dilution); goat anti-FLAG (RRID_Ab: 10000565, 1:500 dilution); and chicken anti-GFP (1:500 dilution). On the following morning, neurons were washed then incubated with Duolink anti-mouse MINUS (DUO 92004) and anti-goat PLUS (DUO92003) probes along with rabbit anti-chicken Alexa Fluor 488 (1:500 dilution) for 1 hour at 37 °C. Probes were ligated at 37 °C for 30 min and signals were amplified at 37 °C for 100 min prior to mounting in ProLong Gold and sealing. Coverslips were imaged within 3 days of PLA labelling. In all cases, imaging was performed using a Zeiss LSM 780 microscope equipped with an airyscan module and using a 60× oil objective lens (Numerical Aperture = 1.40). Z-stacks of 0.38 μm optical slices were collected at 1024 × 1024 resolution. Images were collected using 488 nm excitation/521 nm emission for detecting anti-GFP labelling, and 594 nm excitation/619 nm emission for detecting PLA puncta. Images were analysed using NeuronStudio software (Icahn School of Medicine at Mount Sinai) to determine spine width and morphology; and the Distance Analysis (DiAna) plugin (Gilles et al., 2017) for ImageJ (NIH) to identify PLA puncta. For each neuron, spine or PLA puncta frequency per 10 μm dendrite was calculated across all clearly-resolved segments of the dendritic arbor. Normality of PLA and spine width data was confirmed using Kolmogorov-Smirnov testing prior to unpaired two-tailed Student’s t-tests.

### Measurement of miniature EPSCs

mEPSCs were recorded 3-4 days after transfection from GFP-expressing primary hippomcapal neurons transfected with either pIRES-CaMKIIα WT or AA. Control (GFP negative) neurons were recorded from the same coverslips. Neurons were visualized using an upright microscope (BX51WI; Olympus) equipped with a 20x NA objective (Olympus), fluorescence LED illumination (CoolLED pE-100), and filters (Chroma Technology ET470/40x, 495LP and ET 525/50m) for GFP visualization. The extracellular solution, adjusted to pH 7.3 with NaOH, contained: 145 mM NaCl, 2.5 mM KCl, 2 mM CaCl_2_, 1 mM MgCl_2_, 10 mM glucose, and 10 mM HEPES. To this we added 1 μM TTX, 20 μM D-AP5, and 20 μM SR-95531 to block voltage-gated sodium channels, NMDA receptors, and GABA_A_ receptors, respectively. Whole-cell patch-clamp recordings were made using electrodes pulled from borosilicate glass that had a resistance of 5–7 MΩ when filled with an internal solution containing 140 mM CsCl, 2 mM NaCl, 2 mM MgCl_2_, 0.5 mM CaCl_2_, 2 mM Na_2_ATP, 5 mM EGTA, 0.5 mM Na_2_GTP, 2 mM QX-314 bromide and 10 mM HEPES (adjusted to pH 7.3 with CsOH). Recordings were made from four independent cultures after transfection using calcium phosphate. Currents were recorded at room temperature (22–26 °C) from cells voltage-clamped at −60 mV using an Axopatch 200B amplifier (Molecular Devices). Records were low-pass filtered at 2 kHz and sampled at 20 kHz using a Digidata 1200 interface and pClamp software (Molecular Devices). Series resistance (R_series_) and input capacitance were read directly from the amplifier settings used to minimize the current responses to 5 mV hyperpolarizing voltage steps. R_series_ (6.5–25.0 MΩ) was compensated (40–70 %) and monitored throughout each recording; if a cell showed a >30 % change in R_series_ it was excluded from the analysis. The compensated R_series_ was similar across the three experimental groups (4.9 ± 0.3, 4.8 ± 0.3 and 5.0 ± 0.5 MΩ, for control, WT and AA, respectively). All experiments were performed in an interleaved and blinded manner with unblinding only after completion of the analyses.

mEPSCs were detected using open-source Python software miniML, a deep learning-based detection method trained on a dataset of annotated mEPSCs from cerebellar mossy fiber to granule cell synapses (66). The pretrained cerebellar model was used directly, with a threshold of 0.6 applied to the prediction trace to extract data segments containing mEPSCs. Extracted events were examined individually using NeuroMatic (67) running in Igor Pro (Wavemetrics) and the peak amplitudes and 10-90 % risetimes of single mEPSCs measured. Spurious events were rejected. Overlapping currents and those with clearly notched rising phases, judged to result from closely-timed release events, were included in the calculation of mEPSC frequency but not amplitude. For each recording, an estimate of the baseline current noise was obtained by generating all-point amplitude histograms from 3–5 sections of the record, fitting the most negative current values in each with a single-sided Gaussian and averaging the obtained measures of standard deviation (range 2.3–6.3 pA), which did not differ across groups. The collected measures of individual mEPSC amplitude and 10-90 % risetime were analysed using R (version 4.3.1, the R Foundation for Statistical Computing) and RStudio (version 2024.04.2+764, Posit Software). Measures from untransfected cells from both groups were pooled and compared with those from the two groups of transfected cells. The mEPSC frequency was determined as the total number of mEPSCs detected/record length, and a mean mEPSC waveform was constructed from those events that displayed a monotonic rise and an uncontaminated decay.

## Data availability

Coordinates and structure factors for crystal structures have been deposited with the RCSB Protein Databank with the following accession IDs: 6TS3 (EF34 complex with CaMKIIα regulatory segment peptide); 7B55 (EF34 complex with apo CaMKIIα 1-315); 7B56 (EF34 complex with CaMKIIα 1-315 + AMP-PNP); 7B57 (EF34 complex with CaMKIIα 1-315 + ADP).

## Acknowledgements

This work was supported by BBSRC grants BB/N015274/1 and BB/X008215/1 to MG, a BBSRC studentship to AC, and MRC grant MR/T002506/1 to MF. We are grateful for the support of Nikos Pinotsis in the ISMB Protein Crystallography and Biophysics Centre for assistance with x-ray crystallography and to Ben Tagg for assistance with Python.

## Supplementary Figures

**Figure S1.**
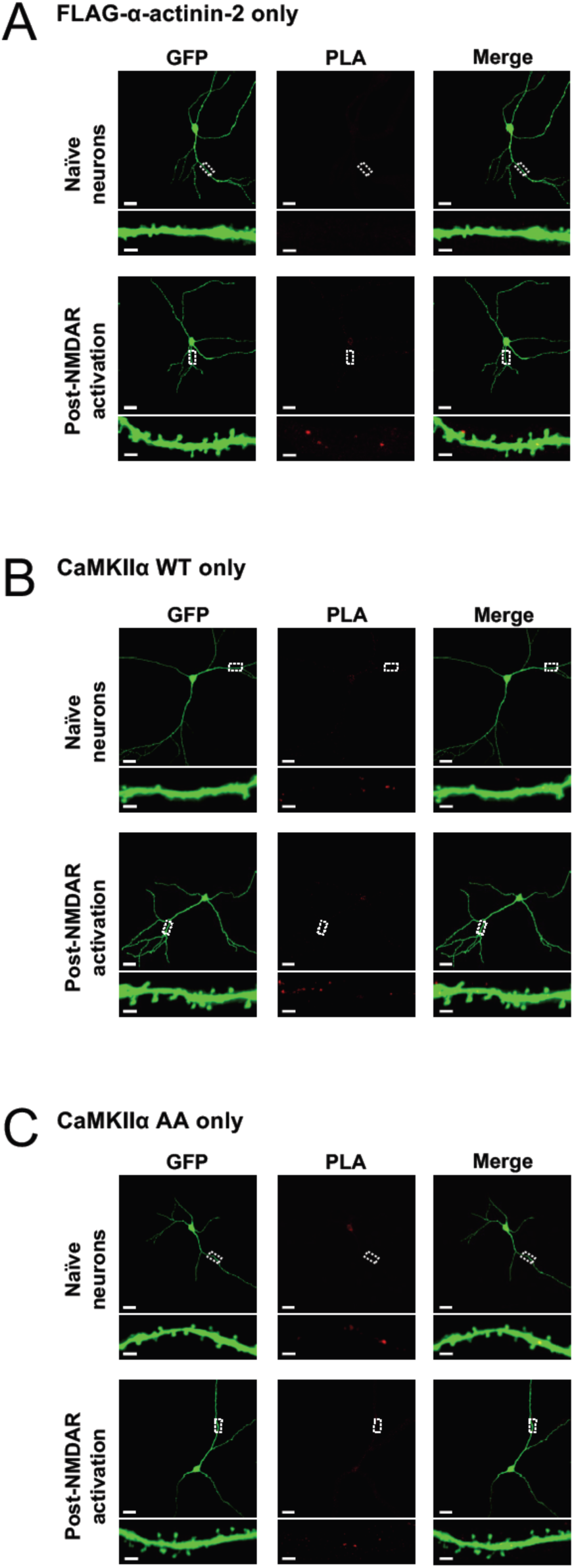
PLA imaging of neurons transfected with single DNA constructs. Each panel shows anti-GFP immunofluorescence (left columns) and anti-FLAG/anti-HA PLA puncta (middle columns) in primary hippocampal neurons either before (top row) or after (bottom row) NMDAR activation. The panels correspond to neurons transfected with pIRES2-GFP constructs expressing either V5-CaMKII⍺ WT (A), V5-CaMKII⍺ T305A/T306A (B), or FLAG-⍺-actinin-2 (C). Scale bars correspond to 20 𝜇m (square panels) and 2 𝜇m (dendrite close-ups).

**Figure S2.**
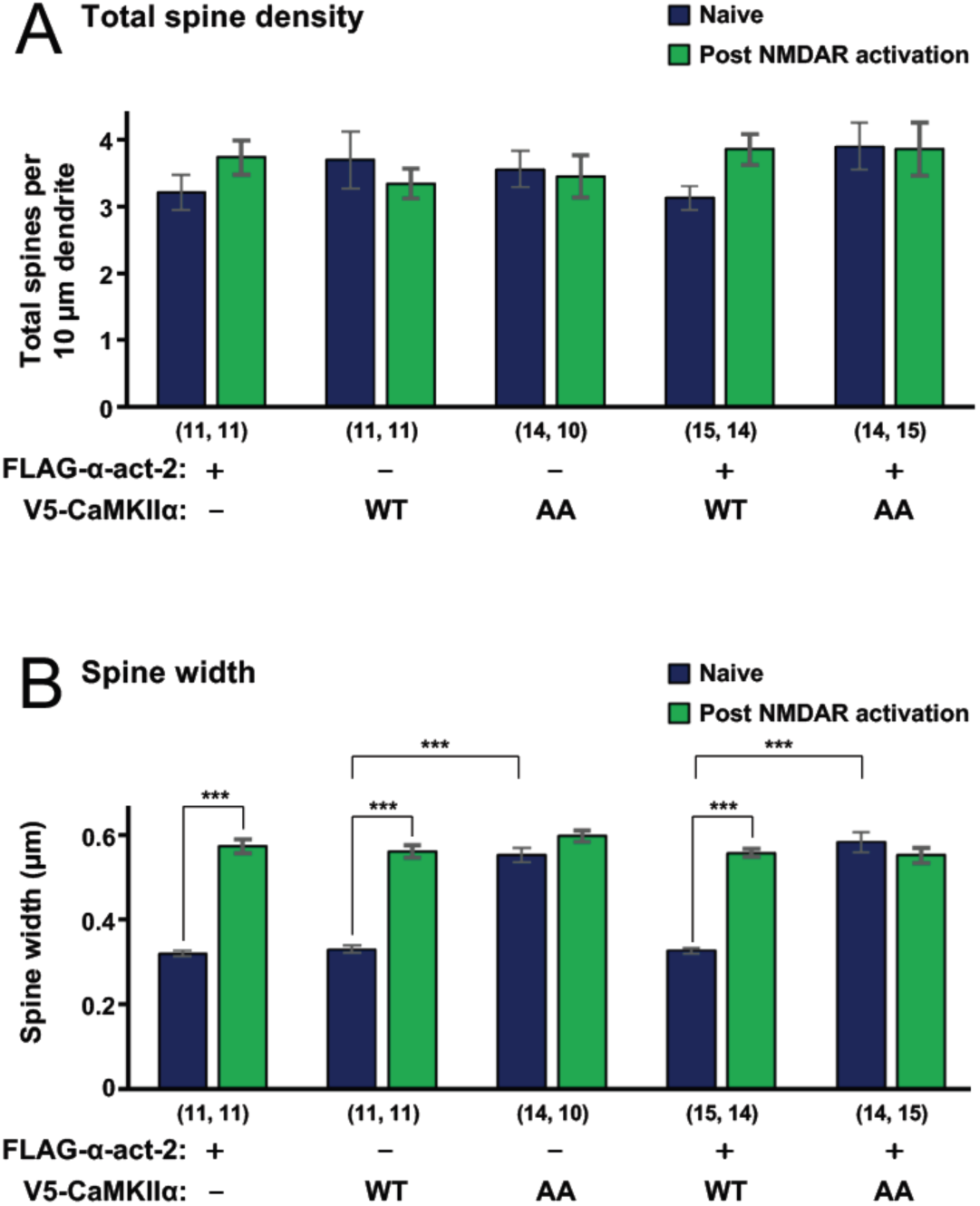
Total spine density and spine width in neurons transfected with actinin and CaMKII variants. Total spine density (A) and spine width (B) in primary hippocampal neurons transfected with different combinations of pIRES-GFP vectors expressing FLAG-⍺-actinin-2 or V5-CaMKII⍺ variants, as indicated. Total spine number per 10 µm dendrite and spine width are shown as mean ± SE before (blue) and after (green) NMDAR activation. The number of neurons analysed for each condition is shown in parentheses. Neurons were imaged deriving from three independent cultures for each condition. In panel B, data were analyzed using unpaired two-tailed Student’s t-tests (\*\*\**P*< 0.001).

**Figure S3.**
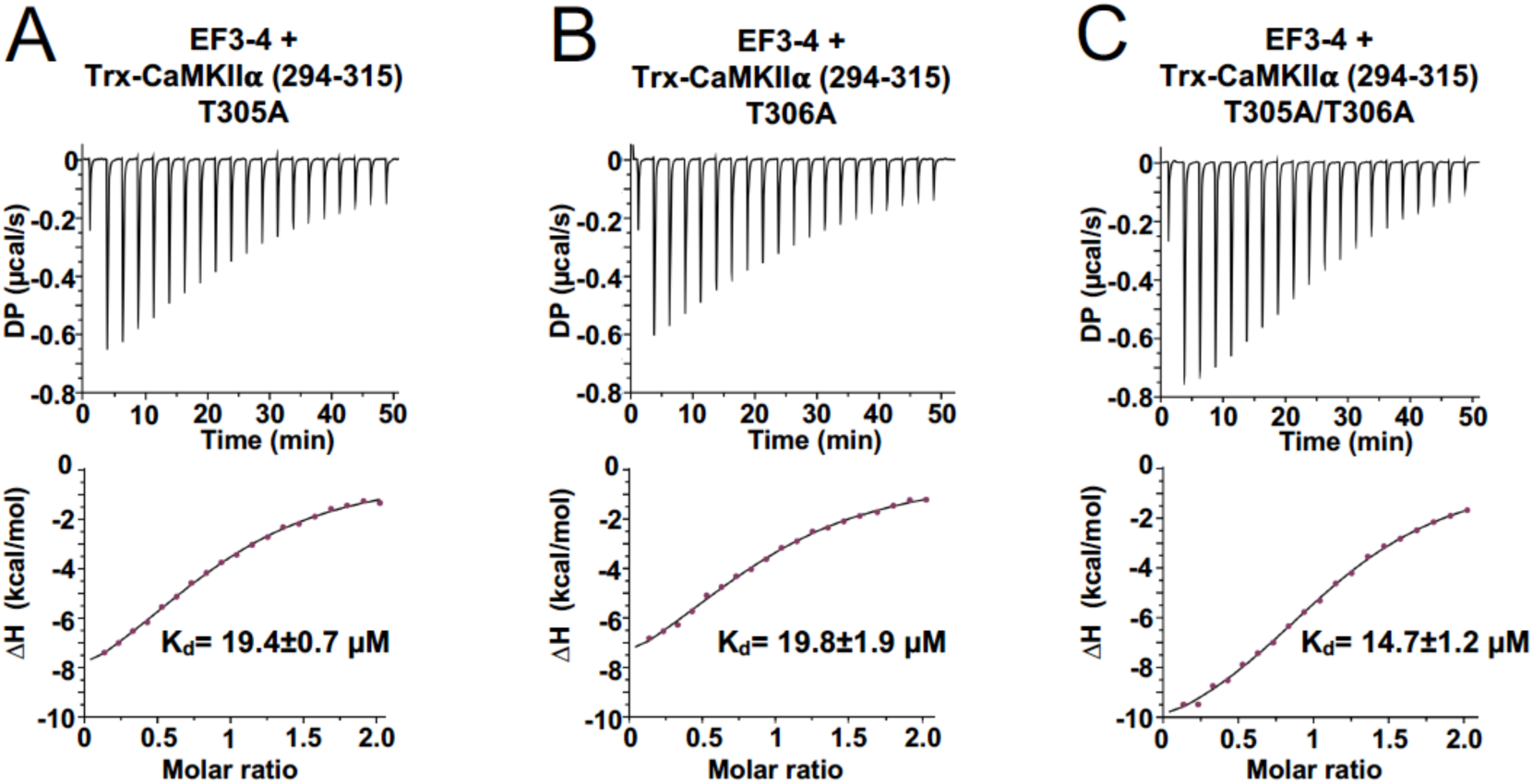
ITC measurements between ⍺-actinin-2 EF3-4 and CaMKII⍺ 294-315 peptides bearing T305A and/or T306A substitutions. Representative isotherms showing the binding of ⍺–actinin-2 EF3-4 to the following CaMKII⍺ 294-315 peptide variants: T305A (A), T306A (B), T305A/T306A (C). The top sub-panels show the raw power output (µcal/s) over time; the bottom sub-panels show the integrated data including a line of best fit to a single site binding model. Stated K_d_ values are averages from experimental replicates.

**Figure S4.**
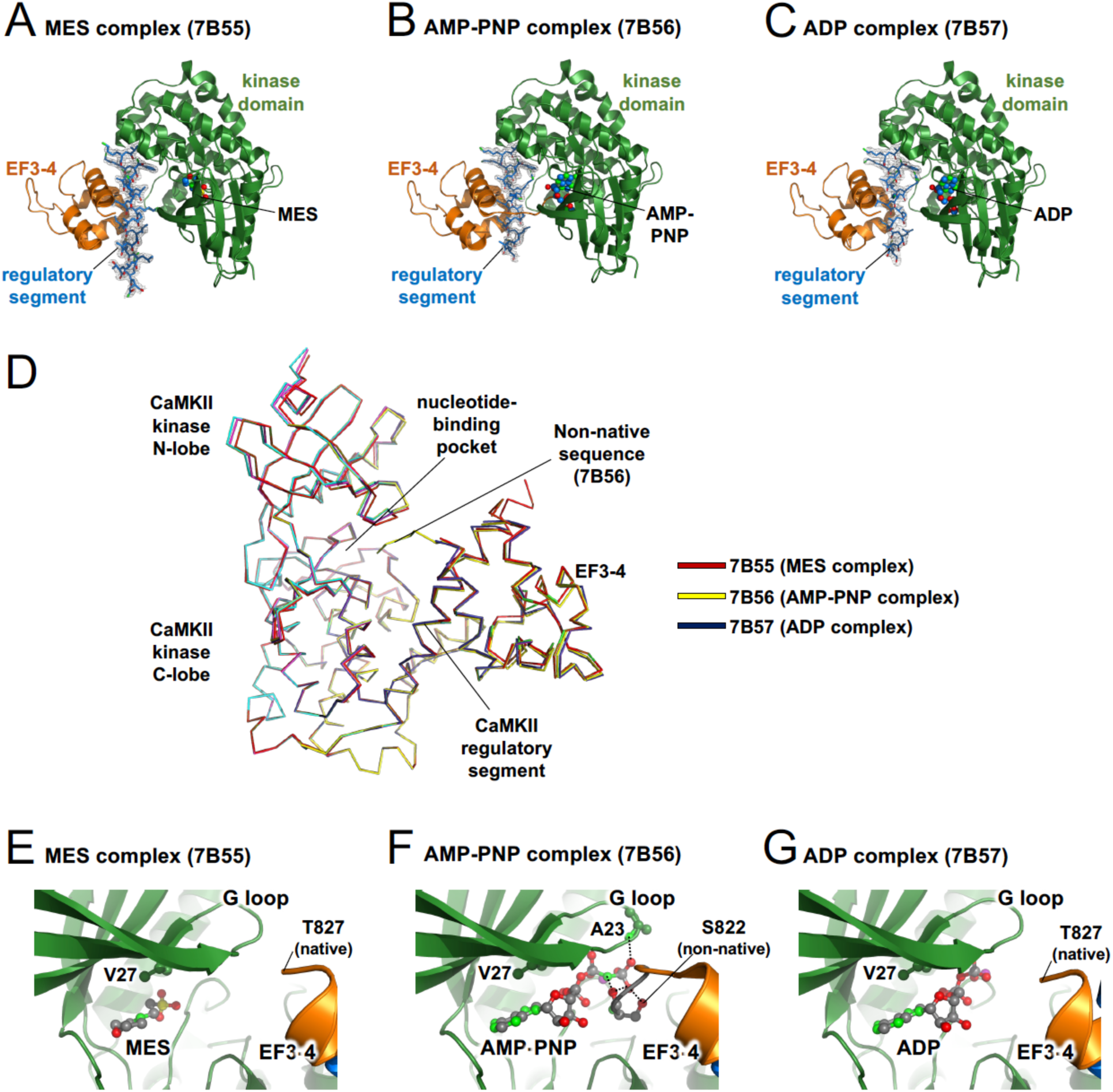
Comparison of crystal structures with different molecules in the nucleotide binding site. Panels A to C show three crystal structures of ⍺-actinin-2 EF3-4 bound to CaMKII⍺ AA that we obtained following crystallisation in buffer containing either no nucleotide (PDB ID 7B55, panel A), Mg^2+^/AMP-PNP (7B56, panel B), or Mg^2+^/ADP (7B57, panel C). The buffer component MES occupies the nucleotide binding site in 7B55 (A). For each structure, the 2F_o_-F_c_ electron density map is shown contoured at 1 𝜎 and clipped within 1.5 Å of the CaMKII𝛼 regulatory segment (blue). (D) Alignment of the three structures with C⍺ positions represented as a ribbon. Panels E to G show close ups of the nucleotide binding sites for structures 7B55 (E), 7B56 (F), and 7B57 (G).

**Figure S5.**
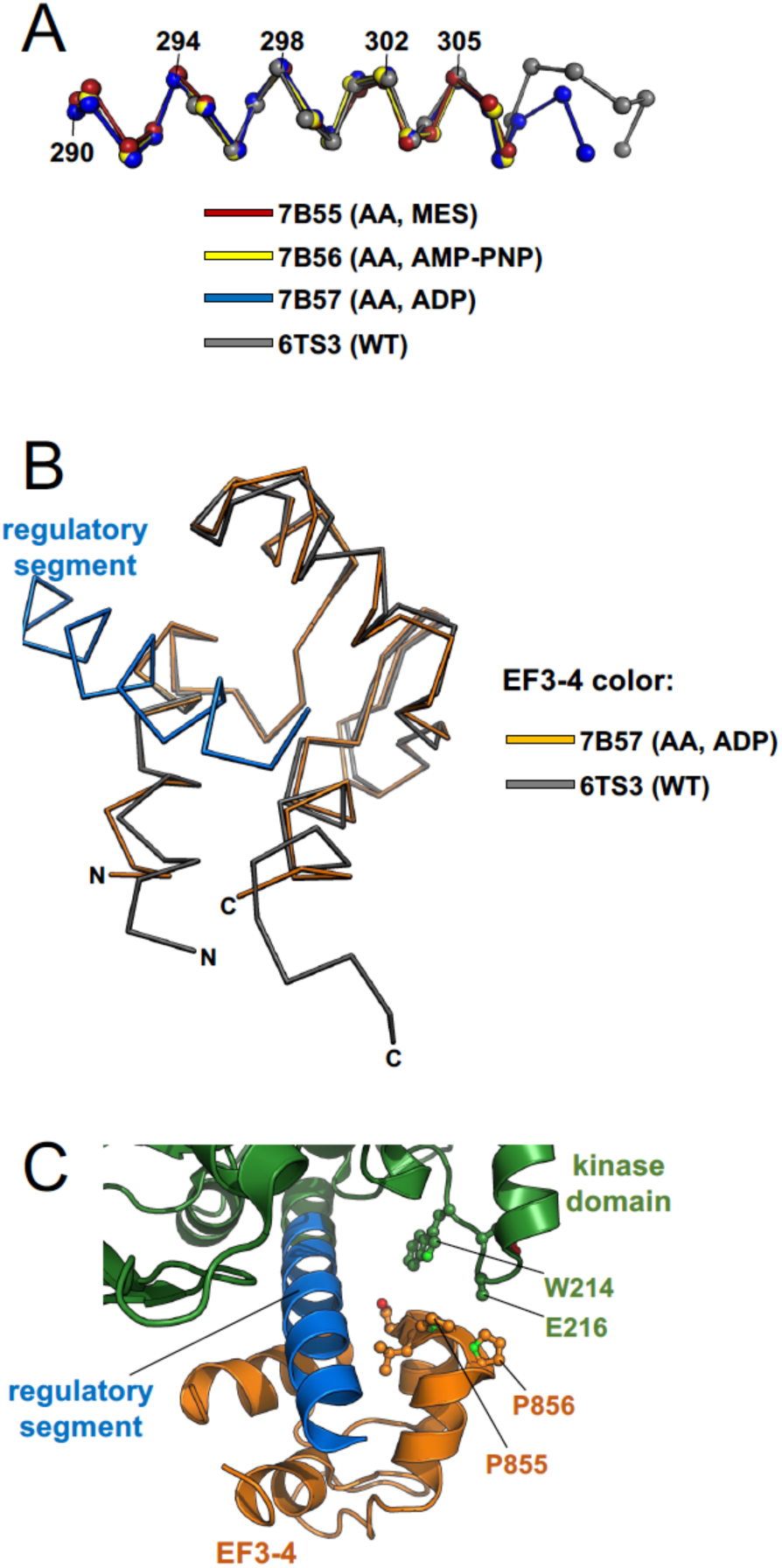
Further structural analysis of CaMKII – ⍺-actinin-2 interactions. (A) Alignment of CaMKII regulatory segment C⍺ positions taken from the three structures determined in this study and from the WT CaMKII⍺ regulatory segment – EF3-4 complex (6TS3). The conformation of the segment is highly similar within 0.6 Å RMSD for all pairwise alignments of positions that are visible across all four structures (294-308). (B) Superposition of EF3-4 taken from the structure in complex with WT CaMKII⍺ regulatory segment (grey) onto the complex with the AA variant, in which EF3-4 is coloured orange. (C) View of the CaMKII⍺ AA – EF3-4 complex (PDB ID 7B57) highlighting the separation of EF3-4 and the CaMKII kinase domain: the two do not engage in any direct interactions.

## Supplementary Tables

**Table S1.** mEPSC measurements in primary hippocampal neurons. . Means of *per-cell* mean measures are presented ± SEM, with the number of cells in parentheses. Each unpaired mean difference (upMD) is given with its 95% bias corrected and accelerated confidence interval (CI) [upper bound, lower bound], calculated form 5000 bootstrap resamples. The *P*-values were determined using a two-sided permutation *t*-test and adjusted for repeated comparison using Holm’s sequential Bonferroni correction. Upper panel: amplitude, 10-90% risetime and frequency measures calculated using all mEPSCs in each cell. Lower panel: amplitude and 10-90% risetime measures calculated after restricting mEPSCs to those whose 10-90% risetime was ≤ 1 ms.

| All mEPSCs | Mean ± SEM (*n*) | upMD [95% CI] | *P* |
| --- | --- | --- | --- |
| **Amplitude (–pA)** |  |  |  |
| Control | 22.0 ± 1.6 (26) | | |
| CaMKIIα WT | 20.2 ± 1.6 (20) | –1.8 [–6.4, 2.2] | 0.93 |
| CaMKIIα AA | 21.7 ± 1.6 (17) | –0.2 [–5.0, 3.9] | 0.93 |
| **Risetime (ms)** |  |  |  |
| Control | 0.98 ± 0.04 (26) | | |
| CaMKIIα WT | 1.01 ± 0.05 (20) | 0.03 [–0.10, 0.15] | 0.69 |
| CaMKIIα AA | 1.06 ± 0.05 (17) | 0.08 [–0.04, 0.20] | 0.43 |
| **Frequency (Hz)** |  |  |  |
| Control | 1.37 ± 0.42 (26) | | |
| CaMKIIα WT | 0.87 ± 0.27 (20) | –0.50 [–1.72, 0.27] | 0.75 |
| CaMKIIα AA | 1.13 ± 0.20 (17) | –0.25 [–1.61, 0.41] | 0.75 |
| **mEPSCs (RT ≤ 1 ms)** | **Mean ± SEM (*n*)** | **upMD [95% CI]** | ***P*** |
| **Amplitude (–pA)** |  |  |  |
| Control | 22.8 ± 1.9 (23) | | |
| CaMKIIα WT | 21.5 ± 2.0 (16) | –1.2 [–7.1, 3.7] | 1.00 |
| CaMKIIα AA | 21.8 ± 1.9 (15) | –1.0 [–6.7, 3.6] | 1.00 |
| **Risetime (ms)** |  |  |  |
| Control | 0.63 ± 0.01 (23) | | |
| CaMKIIα WT | 0.63 ± 0.02 (16) | 0.00 [–0.03, 0.04] | 0.84 |
| CaMKIIα AA | 0.67 ± 0.02 (15) | 0.04 [ 0.00, 0.08] | 0.11 |

**Table S2.** Thermodynamic parameters for interactions between a-actinin EF34 and CaMKII constructs.

| Binding partner A | Binding partner B | Replicates | N | K<sub>d</sub> (μM) | ΔH |
| --- | --- | --- | --- | --- | --- |
| α-actinin-2 EF3-4 | CaMKIIα 294-315 T305A/T306A | 3 | 1.15 ± 0.02 | 14.7 ± 1.2 | −12.5 ± 0.6 |
| α-actinin-2 EF3-4 | CaMKIIα 294-315 T305A | 3 | 0.98 ± 0.03 | 19.4 ± 0.7 | −11.8 ± 0.6 |
| α-actinin-2 EF3-4 | CaMKIIα 294-315 T306A | 3 | 0.91 ± 0.02 | 19.8 ± 1.9 | −10.3 ± 0.5 |
| α-actinin-2 EF3-4 | CaMKIIα 1-315 T305A/T306A | 3 | 0.99 ± 0.4 | 0.154 ± 0.009 | −9.79 ± 0.56 |
| α-actinin-2 EF3-4 | CaMKIIα 1-315 T305A | 2 | ND | ND | ND |
| α-actinin-2 EF3-4 | CaMKIIα 1-315 T306A | 2 | ND | ND | ND |
| α-actinin-2 EF3-4 | CaMKIIα 290-309 WT | 3 | 1.03 ± 0.01 | 3.46 ± 0.08 | −7.02 ± 0.17 |
| α-actinin-2 EF3-4 | CaMKIIα 290-309 T305A/T306A | 3 | 0.99 ± 0.02 | 0.146 ± 0.001 | −9.05 ± 0.21 |

**Table S3.**
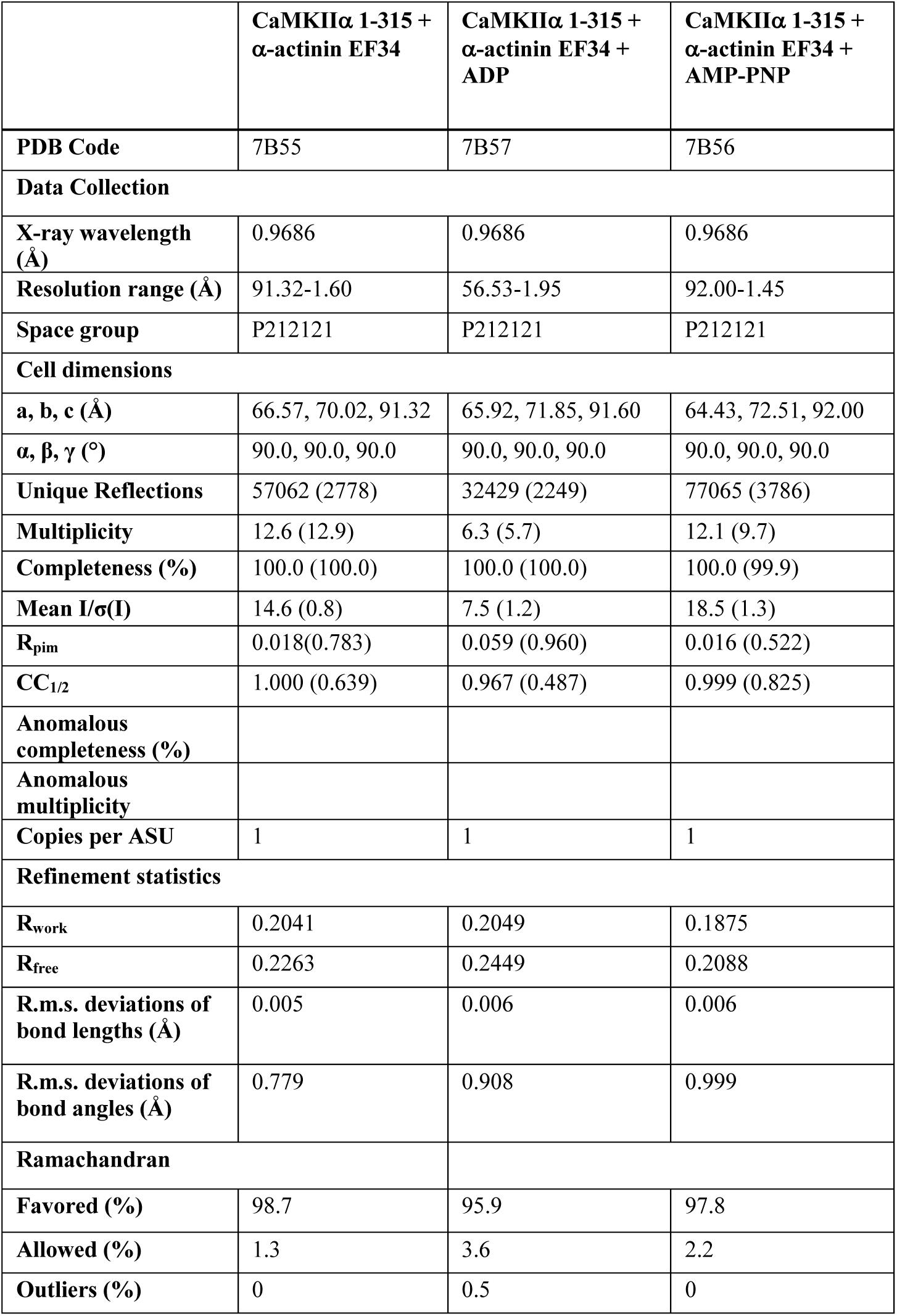
X-ray crystallography data collection and refinement statistics.

**Table S4.**
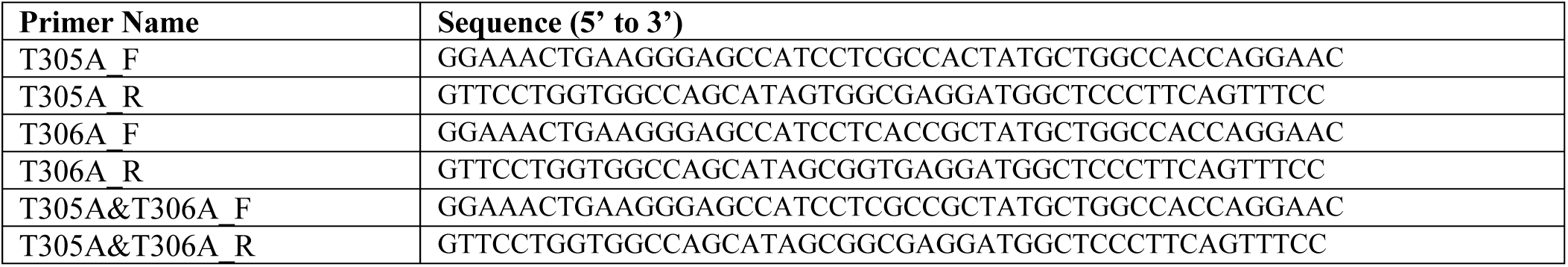
Oligonucleotide primer sequences.

